# Senescent Schwann cells induced by aging and chronic denervation impair axonal regeneration after peripheral nerve injury

**DOI:** 10.1101/2022.12.07.519441

**Authors:** Andrés Fuentes-Flores, Cristian Geronimo-Olvera, David Ñecuñir, Sandip Kumar Patel, Joanna Bons, Megan C. Wright, Daniel Geschwind, Ahmet Hoke, Jose A. Gomez-Sanchez, Birgit Schilling, Judith Campisi, Felipe A. Court

**Affiliations:** Center for integrative Biology, Universidad Mayor, Chile; FONDAP Geroscience Center for Brain Health and Metabolism; Buck Institute for Research on Aging, Novato, USA; Departments of Neurology and Neuroscience, Johns Hopkins School of Medicine, Baltimore, Maryland, USA; Instituto de Investigación Sanitaria y Biomédica de Alicante (ISABIAL), 03010 Alicante, Spain; Instituto de Neurociencias de Alicante, UMH-CSIC, 03550 San Juan de Alicante, Spain; Departments of Neurology, Psychiatry, and Human Genetics, David Geffen School of Medicine, University of California Los Angeles, California, USA

## Abstract

After peripheral nerve injuries, successful axonal growth and functional recovery requires the reprogramming of Schwann cells into a reparative phenotype, a process dependent on the activation of the transcription factor c-Jun. Nevertheless, axonal regeneration is greatly impaired in aged organisms or after chronic denervation leading to important clinical problems. This regenerative failure has been associated to a diminished c-Jun expression by Schwann cells, but whether the inability of these cells to maintain a repair state is associated to the transition into a phenotype inhibitory for axonal growth, has not been evaluated so far. We find that repair Schwann cells transitions into a senescent phenotype, characterized by diminished c-Jun expression and secretion of factor inhibitory for axonal regeneration in both aging and chronic denervation. In both conditions, elimination of senescent Schwann cells by systemic senolytic drug treatment or genetic targeting improves nerve regeneration and functional recovery in aging and chronic denervation, associated with an upregulation of c-Jun expression and a decrease in nerve inflammation. This work provides the first characterization of senescent Schwann cells and their impact over axonal regeneration in aging and chronic denervation, opening new avenues for enhancing regeneration, and functional recovery after peripheral nerve injuries.

## 1. Introduction

The peripheral nervous system (PNS) exhibits an effective regenerative capacity after nerve injury due to a coordinated tissue response. After nerve damage, injured neurons activate an intrinsic growth response and extend over a permissive regenerative environment generated by Schwann cells (SCs), the glial cell component of the PNS. This response to nerve damage is critically dependent on the capacity of SCs to reprogram into a repair phenotype (rSC) which supports axonal regeneration. SCs transition into rSCs is triggered by the injury-induced degeneration of their associated axons and dependent on the activation of the transcription factor c-Jun^1, 2^. SCs reprogramming is characterized by cell proliferation, myelin phagocytosis and the secretion of pro-regenerative factors, including exosomes^2–4^. In addition, rSCs recruit macrophages and generate cell tracks that guide regenerating axons^5^.

Despite the good regenerative capacity of the PNS, especially in laboratory setups, nerve injuries remain a major cause of morbidity in the clinic^6–8^. Less than half of the patients suffering nerve injuries will recover useful function, and up to one-third will experience little or no recovery after appropriate surgical intervention^7–9^. Importantly, previous work has established that age and chronic SC denervation are the major factors affecting peripheral regeneration in laboratory models and humans^10–13^. Age-dependent regeneration deficits has been associated with an enhanced inflammatory response in the nerve and a decrease capacity of SCs to maintain their reparative phenotype^14^. Interestingly, this age-dependent decrease in regeneration is not due to a reduction in the growth capacity of regenerating neurons^10, 11, 15, 16^. In the case of chronic denervation, it has been reported that rSCs gradually lose their ability to secrete pro-regenerative trophic factors such as BDNF or GDNF^13, 17^. Importantly, chronic SC denervation arise not only as a consequence of delayed surgical repair, but also as a result of axons regenerating for months or even years in long human nerves^13, 17, 18^. It has been recently demonstrated that aged and chronically denervated rSCs exhibit diminished c-Jun expression after injury, and forced upregulation of this transcription factor restores axonal regeneration in these two conditions^19^. Even though high c-Jun expression is sufficient to activate axonal regeneration, the phenotype acquired by low-c-Jun expressing SCs has not been clearly defined, nor a possible inhibitory effect they might have over axonal regeneration. Interestingly, it has been shown that SCs invading long acellularized nerve allografts express markers of senescent cells^20, 21^. This raise the question of whether SCs undergo senescence in the context of aging and chronic denervation, and if this senescent SC phenotype is associated with downregulation of c-Jun expression, and the production of inhibitory factor(s) for axonal regeneration and functional recovery.

Senescence is a cellular state induced in response to various stressors, such us irreparable DNA damage, oxidative stress and oncogenic activation, and characterized by an irreversible arrest of the cell cycle and resistance to apoptosis^22–24^. Senescent cells show chromatin instability, increased cellular size, enhanced lysosomal activity, abnormalities of nuclear morphology and a specific senescence-associated secretory phenotype (SASP) composed of pro inflammatory angiogenic and extracellular matrix degrading factors^22, 25, 26^. It has been shown that senescent cells can have both, a beneficial and detrimental role in tissue repair, depending if the exposure to the stressing stimuli is transient or chronic^27, 28^. After acute tissue damage, senescent cells are necessary to inhibit the proliferation of damaged cells and prevent tumor formation through the wound healing process, after which they are cleared by macrophages^27, 29^. In contrast, chronic exposure to stressor agents, stimuli or aging generates chronic accumulation of senescent cells. Chronic senescent cells are inefficiently cleared, contributing to abnormal SASP accumulation in the tissue. This affects neighboring cells in a paracrine fashion, leading to tissue disfunction or age-related pathologies^23, 27, 30, 31^. In this regard, chronically denervated SCs stop proliferating through cell cycle arrest and also show reduced expression of essential pro-regenerative factors such as BDNF, GDNF and NRG1^32^, with increased secretion of proinflammatory cytokines^13, 18, 33^.

Here, we characterized senescent Schwann cells (sSC) and their role over axonal regeneration after peripheral nerve injury. We found that c-Jun-negative SCs accumulate after injury in aged and chronically denervated nerves exhibiting *bona fide* markers of cell senescence. In contrast to the pro-regenerative effect of rSC secreted factors, the senescent SC-associated secretory component strongly inhibits axonal growth of sensory neurons. Importantly, systemic treatment with a senolytic eliminates senescent Schwann cells (sSC) and enhances axonal regeneration in aged and chronically denervated animals. Elimination of sSC downregulates pro-inflammatory factors present in injured nerves from aged animals and after chronic denervation, restoring c-Jun levels in SCs. Taken together, our data demonstrate that sSC have an inhibitory effect over axonal regeneration in aging and chronic denervation. Therefore, elimination of sSC or neutralization of their inhibitory secreted factors, stands out as a novel therapeutic intervention to enhance axonal regeneration and functional recovery after peripheral injuries.

## 2. Results

### 2.1. **Senescent Schwann cells accumulate in peripheral nerves after chronic denervation and in aged animals.**

As up-regulation of c-Jun in rSC after damage is essential for effective axonal regeneration, we first evaluated the dynamics of c-Jun expression in SCs after sciatic nerve cut under two conditions in which regeneration is impaired, aging and chronic denervation. Consistent with previous studies^19^, we detected a significant increase in the expression of c-Jun in denervated SCs from adult (2-4 months old) mice 7 days post-injury (dpi) compared to undamaged control nerves. By contrast, in aged animals (20-22 months old), denervated SCs fail to up-regulate c-Jun to levels comparable to adult denervated SCs (Figures1A and S1B-C). After chronic SC denervation (21 and 42 dpi), c-Jun levels drop dramatically in both aged and adult nerves (Figure 1A and S1B- and S1C). We then studied axonal regeneration by using a model of nerve repair in which the tibial branch was first transected and the distal stump sutured to the nearest muscle to prevent reconnection. After 12 or 42 dpi, the distal tibial nerve was detached from the muscle and reconnected to the freshly transected peroneal common nerve, and axonal regeneration was evaluated after 7 days (Figure S2). In both chronic denervation (42 dpi) and aging (12 dpi), axonal regeneration was strongly inhibited compared to adult animals reconnected at 12 dpi (Figures 1B-1C).

**Figure 1.**
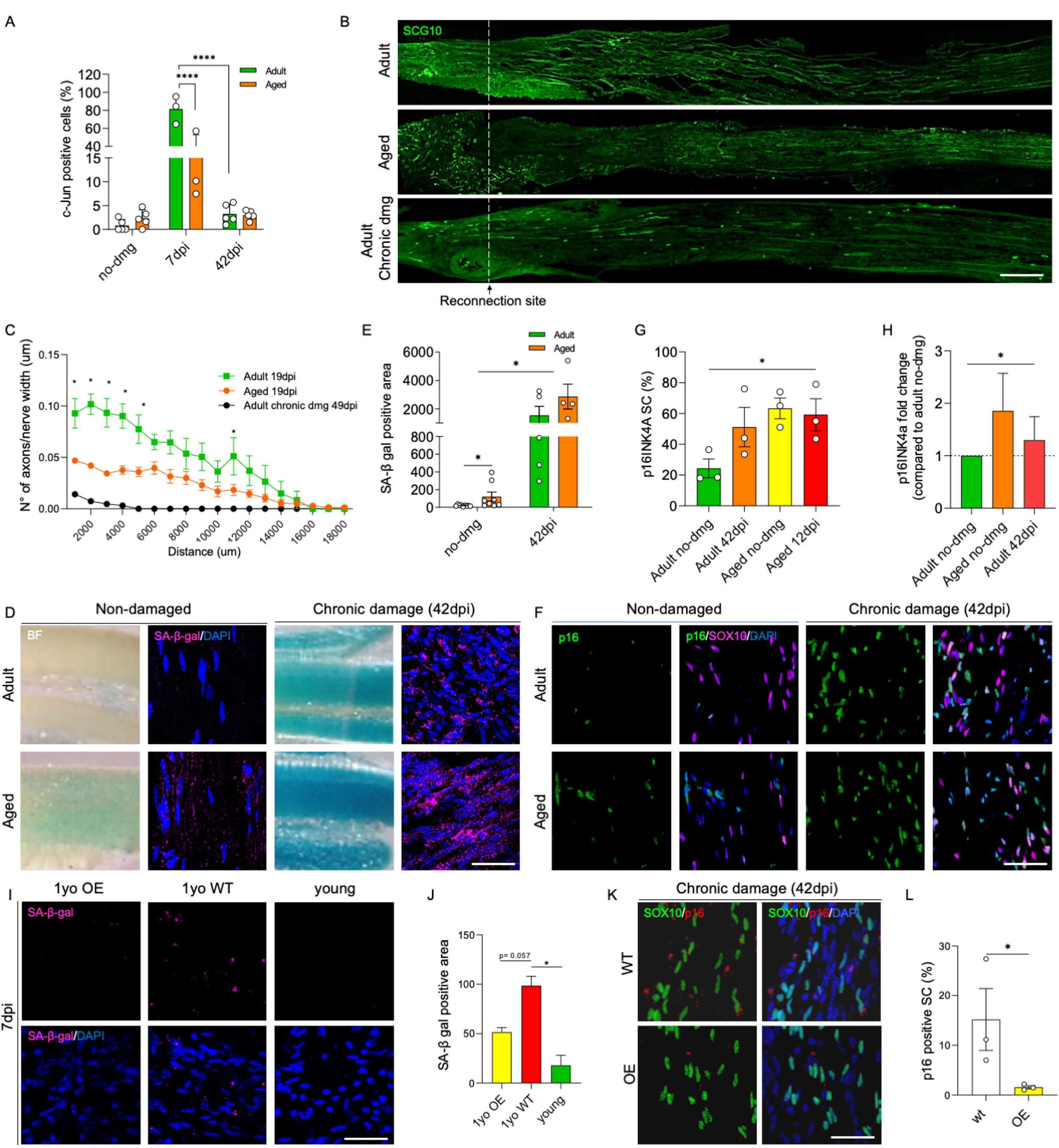
Aged and chronically denervated sciatic nerves show reduced regeneration and higher senescent cell accumulation. (A) c-Jun activation in the nucleus of SCs on longitudinal cryostat sections of adult and aged mice sciatic nerves in non-damaged animals (ctrl), and sciatic nerves distal to the nerve cut at 7 and 42 dpi (n=3-5 animals per group, p<0.05 by Student’s t-test compared between conditions; error bars indicate S.D.). (B) Representative IF images of sciatic nerves 7 days after reconnection surgery in aged, adult (reconnected at 12 days after damage) and chronically denervated (reconnected at 42 days after damage) (SCG10, green). Scale bar, 500 μM. (C) Axonal density and distance comparison 7 days after reconnection surgery in aged, adult (reconnected at 12 days after damage) and chronically denervated nerves (reconnected at 42 days after damage) animals. The dataset used for Figure 1B and C corresponds to control experiments of the experiment shown in Figure 5A-C. (D, E) Brightfield and fluorescence confocal acquisition of β-galactosidase activity on longitudinal sections of non-injured nerves from adult and aged animals, and nerves distal to the nerve cut at 42 dpi in adult and aged animals after b-galactosidase assay. (β-gal, magenta; DAPI, blue). (n=5 animals per group, p<0.05 by Student’s t-test compared between conditions; error bars indicate S.D.) Scale bar, 100 μM. Complete nerves in brightfield are shown in Figure S1a. (F-H) qRT-PCR and fluorescence quantification of p16INK4a in contralateral non-injured nerves and chronically denervated (42 dpi) sciatic nerves of adult and aged animals. Scale bar, 50 μM. (I-J) Fluorescence confocal acquisition of β-galactosidase activity on longitudinal sections of injured sciatic nerves distal to the nerve cut at 7 dpi of adult WT and c-Jun OE animals after b-galactosidase assay. Scale bar, 50 μM. (β-gal, magenta; DAPI, blue). (K-L) qRT-PCR and fluorescence quantification of p16INK4a in sciatic nerves distal to the nerve cut at 42 dpi (chronically denervated) from adult WT and c-Jun OE animals. Scale bar, 50 μM. (n=5 animals per group, p<0.05 by Student’s t-test compared between conditions; error bars indicate S.D.).

As chronic proliferative phenotypes and aging has been associated to senescent cell accumulation^31^, we assessed if the impairment in axonal regeneration in aging and chronic denervation were correlated with accumulation of senescent cells after nerve injury. To this end, we used senescence markers in aged or chronically denervated nerves *in vivo*. SA-β-galactosidase (β-gal) expression, a canonical senescent-cell marker, was increased in uninjured aged nerves compared to adult animals (Figure 1D and 1E). When β-gal was evaluated in the distal nerve 7 dpi, adult and aged animals showed equivalent β-gal activity, which increased with chronic denervation at 42 dpi (Figure 1D, 1E and S1A). As senescence characterization requires the combination of several markers^34^, we measured the expression and localization of the cyclin-dependent kinase (CDK) inhibitor p16INK4a (cdkn2a), critical for the cell cycle arrest and senescence development^35^ on SOX10-positive SCs. Importantly, the percentage of p16INK4a-positive SCs were strongly increased in injured aged nerves in comparison to uninjured adult ones (Figure 1F-1G). The expression of p16INK4a in SCs was also significantly increased in chronically denervated nerves of both adult and aged animals (Figure 1H).

As a decrease in c-Jun expression is associated to impaired nerve regeneration, we next asked whether this decrease was associated to the appearance of senescent SCs. Importantly, previous reports have shown an inhibitory feedback loop between p16INK4a and c-Jun^36, 37^. To determine the functional significance of this, we studied the SC response in mice in which c-Jun levels are overexpressed only in SCs (*Mpz*Cre+;R26c-Junstopff/+ mice, referred to as c-Jun OE mice). Importantly, c-Jun overexpression in this model restores regeneration after injury in aged animals and after chronic denervation^38^. Suggesting a c-Jun dependent control of SC senescence, a robust decrease in SA-β-gal activity after nerve transection was observed in 1-year old c-Jun OE mice compared to wildtype animals of the same age (Figure 1I, 1J). Additionally, a significant decrease in p16INK4a-positive SCs after chronic denervation was also observed in the c-Jun OE mice compared to control mice (Figure 1K, 1L). Taken together, our results indicate an increased accumulation of senescent SCs in both aged and chronically denervated neuronal tracts, which correlates with the decrease in c-Jun expression and regenerative capabilities.

### 2.2. Injury-induced nerve inflammation is enhanced in aged and chronically denervated animals

As senescent cells express a secretory phenotype (SASP) enriched in proinflammatory cytokines^31, 39^ and previous studies have shown an increase in inflammation in aged and chronically denervated nerves^14^, we studied if senescent cells could be associated to this inflammatory reaction in aged and chronically denervated nerves. We first evaluated the increase in cytokines from aged (12 dpi) and chronically denervated nerves (42 dpi), both conditions evaluated in our previous experiment (Figure 1C), compared to a nerve severed for 7 days (7 dpi), which our data shows have the higher c-Jun expression in SCs, therefore regenerative potential. From a total of 111 analytes assessed using a cytokine array, 99 and 101 were increased at least 2-fold in chronic denervation and aging, respectively, compared to adults at 7 dpi (Figure 2A, Table 1 and Table 2). Of these analytes, 17 were previously described as SASP components, leaving 82 (chronic denervation) and 84 (aging) secreted proteins as new molecular components of aged and chronically denervated nerves. We next performed a transcriptomic analysis to search for the expression of SASP-associated genes in nerves chronically denervated. Our analysis revealed that 250 of 1646 differentially up regulated genes ranging from acute (1 day) to chronically denervated nerves (180 days), were previously reported as SASP related genes (Figure S3A-B and Table 3). This further validated the increase in SASP components at the transcriptional level in chronically denervated conditions. Interestingly, the proportion of unique SASP components increases with the time of denervation (Figure S3C). These unique SASP genes range from 10% to 30% of the total SASP as denervation progress through time, defining a unique SASP signature for chronic denervation (Figure S3B-D). Taken together, our data complement previous results on nerve inflammation in aging and chronic denervation, identifying novel molecules upregulated in these two conditions, and providing evidence for development of an inflammatory condition in the nerve. Importantly, the identification of an important number of SASP-associated proteins and transcripts strongly suggests that the increase in nerve inflammation could be closely related to the increase in cell senescence after nerve damage in aging and chronic denervation.

**Figure 2.**
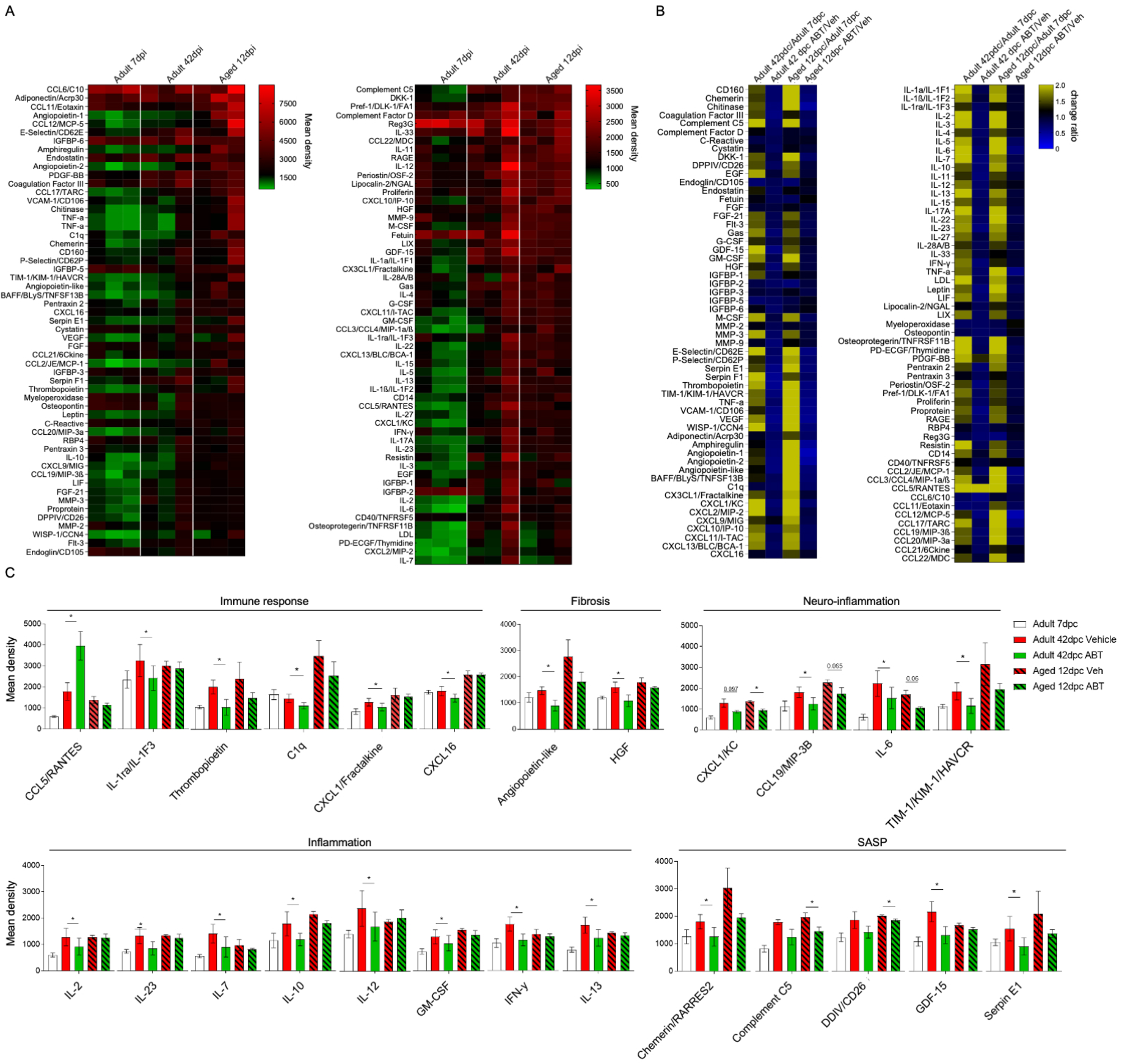
Elimination of senescent SC in aged and chronically denervated nerves alters cytokine profiling *in vivo*. (A) Expression analysis of 111 different cytokines in the context of aging or chronic denervation. Comparison between chronic denervation (Adult 42 dpi) or aging (Aged 12 dpi). (B) Fold change comparison of 111 different cytokines in the context of aging or chronic denervation after ABT treatment. Comparison between chronic denervation (Adult 42 dpi) or aging (Aged 12dpi) conditions in the context of ABT treatment, vehicle treatment or acute regenerative response (Adult 7 dpi). (C) Expression analysis of 111 different cytokines in the context of aging or chronic denervation. Comparison between chronic denervation (Adult 42 dpi) or aging (Aged 12dpi) conditions in the context of ABT treatment, vehicle treatment or acute regenerative response (Adult 7 dpi). Data are expressed as mean density (n=3 animals per group, p<0.05 by Student’s t-test compared between conditions; error bars indicate SEM.).

### 2.3. Senescent Schwann cells are inhibitory for axonal growth of sensory neurons *in vitro*

To establish the effect of senescent Schwann cells on axonal growth we first used an *in vitro* model. rSCs were treated with the anti-carcinogenic drug doxorubicin, a widely used senescence inducer^31^. After doxorubicin treatment for 9 days, more than 90% of treated SCs were positive for SA-β-gal activity (Figure S4A, B). SCs treated with doxorubicin show significant translocation of HMGB1 from nucleus towards the cytosol, increased p-γH2AX foci, and increased p21^CIP1^ positive nuclei, compared to rSC (Figure S4A-G). In addition, lamin B1 expression was increased, together with an increase in lamin B1-positive nuclear invaginations (Figure S4A, H, I). Finally, doxorubicin-treated SC show an increase in p16INK4a expression compared to rSC (Figure S4J). Importantly these senescence-induced SC (siSC) retain their SC phenotype, assessed by the expression of S100 and SOX10 (Figure S4A and 3A). Surprisingly, induction of a senescent phenotype decreases c-Jun expression in siSC compared to rSC (Figure 3A, 3B).

**Figure 3.**
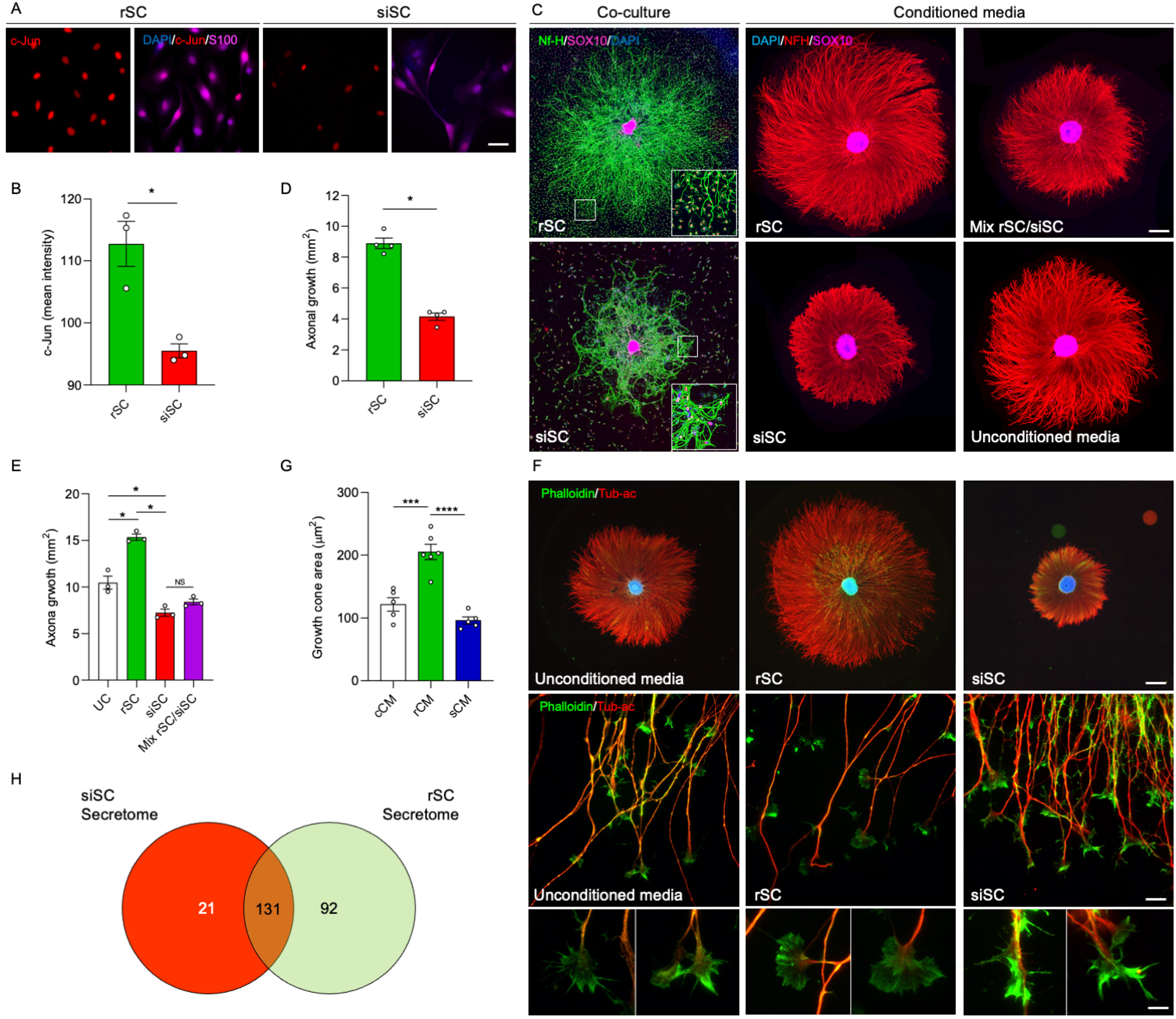
siSCs impairs c-Jun expression and neurite outgrowth *in vitro*. (A-C) Representative IF images (A), graph comparison (B) of c-jun positive SC in the nucleus area between non-senescent and sSCs SCs after doxorubicin treatment (S100, green; c-jun, red; DAPI, blue). Scale bar, 100 μM. (C-E) Representative IF images (C) and graph (D-E) comparison between DRG re-aggregates after 72 hrs of co-culture with rSCs or siSCs and exposed only to conditioned media from rSC or sSCs respectively. (F-G) Representative IF images (F) and graph (G) comparison between DRG explants growth cone area after 72 hrs of exposure to conditioned media from rSC or sSCs respectively (Acetylated tubulin, red; Phalloidin, green; DAPI, blue). (H) Secretome analysis of conditioned media from siSC and rSC compared with SASP-ATLAS database. (n=3-4 re-aggregates per group; * p<0.05 by Student’s t-test compared between conditions; error bars indicate S.D.). Scale bars, C (co-culture), 500 μM, C (conditioned media), 1000 μM; F first row, 1000 μM; F second row, 100 μM; F third row, 50 μM.

We next used siSC to establish their effect on axonal growth. We first cultured explants of sensory neurons over monolayers of rSC or siSC, measuring axonal growth after 72 hrs (Figure S5A). Importantly, siSC have a strong inhibitory effect over axonal growth compared to neurons co-cultured with rSC (Figure 3C and 3D). As the effect of senescent cells is usually associated with their secretory phenotype^39^, we evaluated the effect of the secreted component of SC on axonal growth. We first performed proteomic analysis of the rSC and siSC secretome. Our results indicate that a total of 21 proteins are exclusively expressed in siSC (Figure 3H and Table S1), all of them reported as SASP components by the SASP-Atlas identification tool^40^. We next treated sensory neuron explants with conditioned media (CM) from rSC, siSC and a mix of both (rSC/siSC) and axonal growth was measured 72 hrs after CM treatment (Figure S5B). Conditioned media from rSC enhances axonal growth compared to unconditioned media (UM). By contrast, conditioned media from siSC strongly inhibits axonal growth (Figure 3C, 3E), and was even able to overcome the pro-regenerative effects of rSC media, with levels of inhibition comparable to siSC media alone (Figure 3C, 3E).

As growth cone dynamics is tightly coupled to axonal growth, we analyzed these structures after CM treatment. After treatment with CM derived from rSC, growth cones adopted a lamellipodium-like structure^41, 42^, which was quantitatively different from the more filopodia-like shape exhibited by untreated neurons when the growth cone area was measured (Figure 3F, 3G). In contrast, CM derived from siSC led to a collapse of the whole growth cone, involving retraction of the filopodium and lamellipodium (Figure 3F, 3G).

Altogether these data demonstrate that secreted molecules from siSC, enriched in SASP-components, have a negative effect on axonal growth, and modify structural characteristics at the level of the growth cone.

### 2.4. Senescent cell elimination restores c-Jun levels in SCs, and injury-induced nerve inflammation in aged and chronically denervated animals

Having established that siSC strongly inhibit axonal growth *in vitro*, we then assessed the impact of senescent cells over injury-induced nerve changes *in vivo*. To this end, we eliminated senescent cells using the senolytic drug ABT-263, a specific inhibitor of the anti-apoptotic proteins BCL-2 and BCL-xL^43^. After systemic senolytic treatment, a clear reduction in β-gal staining was observed in aged and chronically denervated nerves compared with vehicle-treated animals (Figure 4A-3B). Additionally, the number of SCs positive for the senescence markers p16INK4a and y-H2AX was significantly reduced after the treatment with ABT-263 (Figure 4C-4F). Surprisingly, the number of c-Jun-positive SCs were significantly increased after senolysis in aged and chronically denervated nerves (Figure 4G, 4H), to expression levels comparable to adult nerves at 7 dpi (Figure 1A and 4G, 4H).

**Figure 4.**
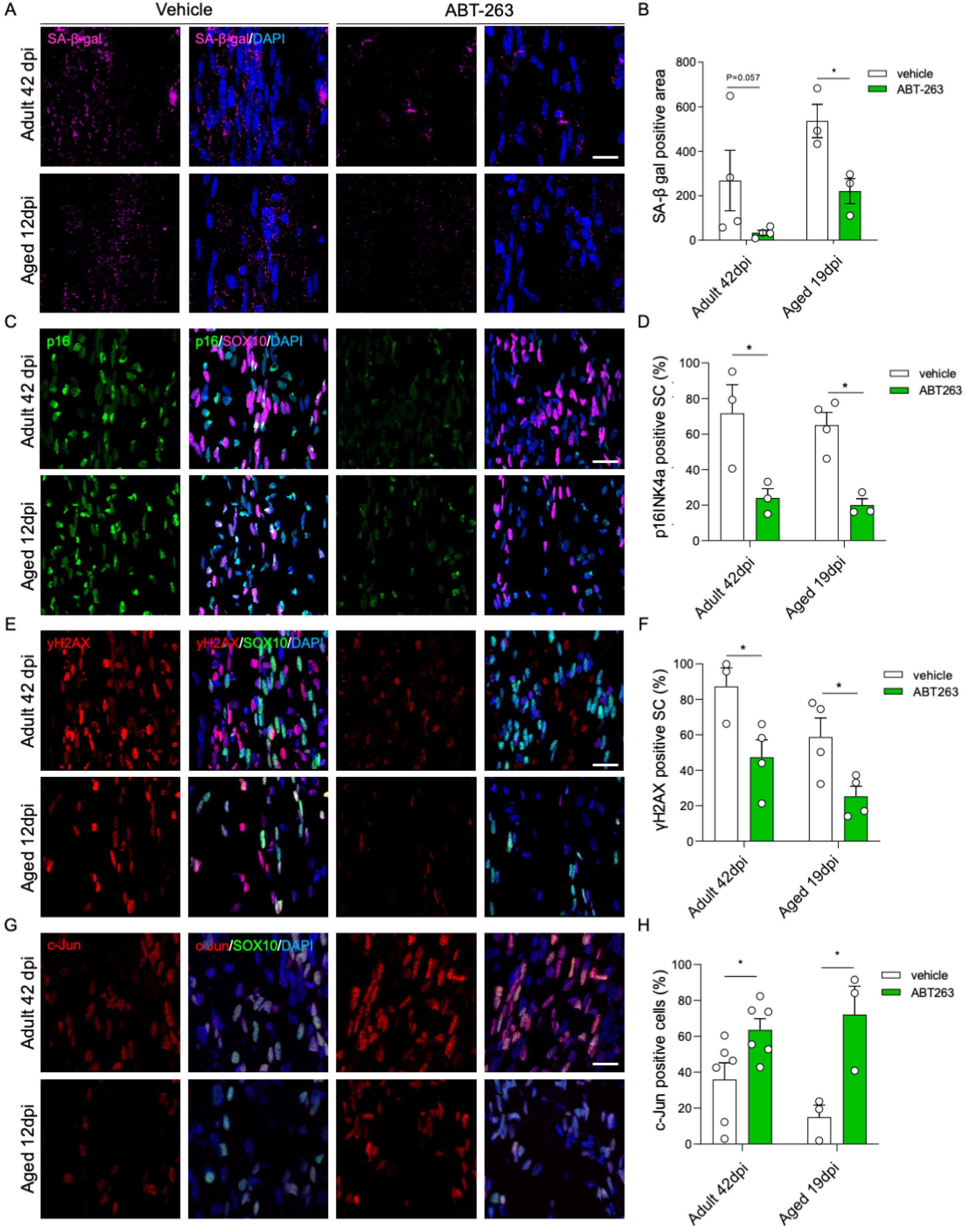
ABT-263 treatment reduces the expression of senescence markers and increases c-Jun translocation in SCs. (A-B) Confocal microscopy and graph comparison of β-galactosidase activity in adult chronically denervated (42 dpi) or aged (12 dpi) animals treated with ABT-263 compared with vehicle treatment, (β-gal: magenta, DAPI: blue). (n=3-5 animals per group, p<0.05 by Student’s t-test compared between conditions; error bars indicate S.D. Scale bar, 100 μM. (C-H) Representative IF confocal images and graph comparison of p16INK4a (green), γ-H2AX (red) and c-Jun (red) positive SCs (SOX10, magenta or green) on longitudinal cryostat sections of adult chronically denervated (42 dpi) or aged (12 dpi) animals, treated with ABT-263 compared with vehicle treatment. (DAPI= blue) (n=3-5 animals per group, p<0.05 by Student’s t-test compared between conditions; error bars indicate S.D.). Scale bars, 50 μM.

We then used the senolytic drug ABT-263 to test the contribution of senescent cells in the inflammatory reaction of aged and chronically denervated nerves. Interestingly, 25 out of 111 proteins that were up-regulated in both aging and chronic denervation (adult 42dpi/aged 12dpi vs adult 7dpi), were significantly decreased after ABT-263 treatment in either one or both conditions (aging and/or chronic denervation, Figure 2B, Table 1 and Table 2). Interestingly, these factors belong to gene ontology (GO) annotations closely related to cell senescence, including immune response, inflammation, fibrosis, neuroinflammation and SASP^35, 44^. Of the 25 identified factors, 5 were previously described as SASP factors, including C5, DPP4, IGFBP-3, Serpin E1, Chemerin/RARRES2, and GDF15^40^, which in both conditions shown a significant decrease after treatment with ABT-263 (Figure 2B). Some of the identified factors, such as CXCL1, CCL19, IL-6 and TIM-1, are significantly decreased in both aging and chronic denervation (Figure 2C), and are tightly related to neuroinflammation upon nerve injury^14, 45–48^. Particularly, CXCL1 is reported to inhibit axonal outgrowth of sensory neurons^45^. These results confirm the elevation of proinflammatory factors previously described in aged nerves after injury and chronically denervated ones and identify several new cytokines that are upregulated in these conditions. Importantly, our data demonstrate that this proinflammatory reaction is associated with a senescent cell population that appears in injured nerves from aged animals and after chronic denervation.

### 2.5. Elimination of senescent cells improves axonal regeneration and functional recovery after nerve injury in aging and chronic denervation

As senescent cell elimination leads to increased c-Jun levels and dampens nerve inflammation in injured nerves from aged animals and after chronic denervation, we assessed the impact of the senolytic ABT-263 on axonal regeneration after nerve repair (Figure S2A-C). As our results demonstrate a basal accumulation of senescent cells in aged nerves before nerve injury (Figure 1D-1H), we treated aged nerves with senolytic immediately after transection, aiming to eliminate resident senescent cells (Figure S2C). 12 days after senolytic treatment, we performed nerve repair surgery and analyzed regeneration after 7 days (Figure S2A, C). Notably, treatment with ABT-263 strongly increase axonal regeneration compared to vehicle-treated animals (Figure 5A-5B), suggesting that resident senescent cells within aged nerves inhibit axonal regeneration.

**Figure 5.**
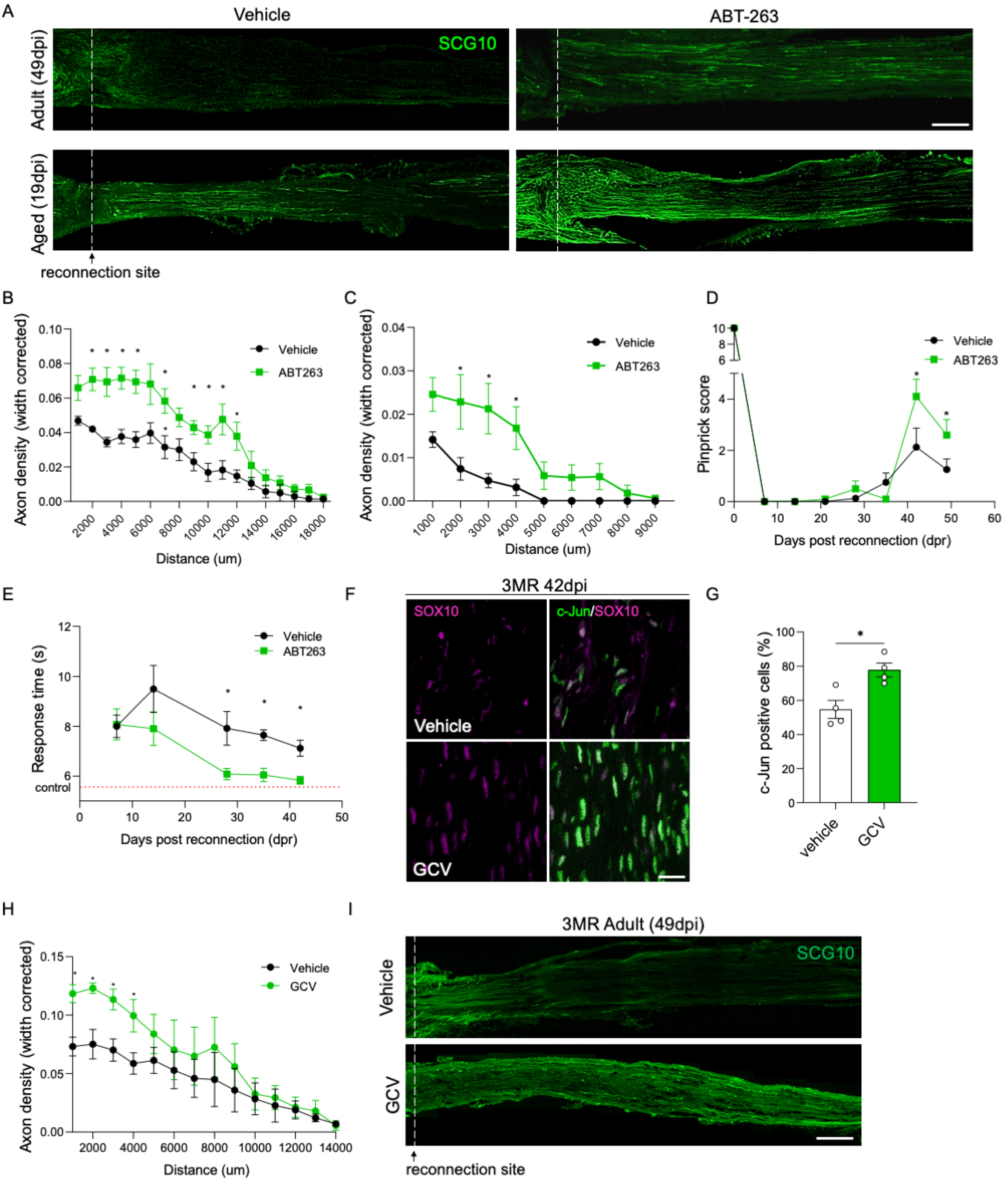
ABT-263 treatment increases axonal regeneration and c-jun expression in SC in aging or chronically denervated sciatic nerves. (A-C) Axonal density and distance comparison, 7 days after reconnection surgery (49 dpi) in adult or aged (19 dpi) animals treated with ABT-263 or vehicle and IF of reconnected nerves (SCG10, green). (n=3-5 animals per group, p<0.05 by Student’s t-test compared between conditions; error bars indicate S.D.). Scale bar, 400 μM. The dataset used for the vehicle controls of Figure 5A-C was used for the experiment shown in Figure 1B and C. (D-E) Mechanical (Pinprick) and thermal (Hargreaves) and functional analysis measured in the injured hindpaw of animals treated with ABT or vehicle solution up to 48 days post reconnection (dpr). (n=8-10 animals per group, p<0.05 by Student’s t-test compared between conditions; error bars indicate S.D.). Response time to thermal stimuli in uninjured animals is shown as a red dotted line in E. (F-G) Representative IF confocal images and graph comparison of c-Jun (red) positive SCs (SOX10, magenta) on longitudinal cryostat sections of adult 3MR chronically denervated (42 dpi) animals, treated with GCV compared with vehicle treatment. (DAPI, blue) (n=3-5 animals per group, p<0.05 by Student’s t-test compared between conditions; error bars indicate S.D.). Scale bar, 50 μM (H-I) Axonal density and distance comparison, 7 days after reconnection surgery (49 dpi) in adult 3MR animals treated with GCV or vehicle and IF of reconnected nerves (SCG10, green). (n=3-5 animals per group, p<0.05 by Student’s t-test compared between conditions; error bars indicate S.D.). Scale bar, 400 μM.

To test the effect of senescent cell elimination over regeneration in chronically denervated nerves, ABT-263 treatment was applied to adult mice 28 days post injury, and after 12 days, nerves were reconnected (Figure S2A, B). Impressively, ABT-263 treatment increases regenerating axons by 2-fold compared to vehicle-treated animals, and almost double the distance of the regenerating front (Figure 5A-5C). To establish if the improvement on axonal regeneration after ABT-263 was associated to a better functional outcome, we assessed recovery of nociceptive sensitivity after reconnection of chronically denervated nerves. Light touch and temperature sensitivity in the skin territory innervated by the tibial nerve was assessed by the pinprick and Hargreaves tests, respectively. Importantly, when compared to vehicle-treated animals ABT-263 treatment led to a better functional recovery of touch and temperature sensitivity at 6- and 4-weeks post reconnection, respectively (Figure 5D-5E).

To further confirm the role of senescent cells in the inhibition of axonal regeneration, we use the p16-3MR transgenic mice to genetically eliminate senescent cells. Senescent cells in p16-3MR mice express the herpes simplex virus 1 thymidine kinase (HSV-TK), thereby sensitizing p16Ink4a-expressing cells to ganciclovir (GCV)^29^. After denervation of p16-3MR mice for 28 days (Figure S2A), we applied daily injections of GCV for 5 days, and after 7 days nerves were reconnected (Figure S2D). Importantly, elimination of senescent cells in p16-3MR mice leads to an increase in c-Jun expression in chronically denervated nerves after GCV treatment (Figure 5F-5G), as it was found after senolysis with ABT-263. Assessment of axonal regeneration reveals an increase in regenerating axons of GCV treated p16-3MR mice compared to vehicle treated ones (Figure 5H-5I). These results confirm that the selective removal of senescent SCs by both pharmacological and genetic means improves both regeneration and functional recovery after chronic denervation (Figure 5A-5I).

## 3. Discussion

For the past two decades it has been recognized that the pro-regenerative effect of SCs over injured axons is reduced in the context of aging and chronic denervation ^18, 49^. Here, we demonstrated for the first-time the presence and functional role of senescent SCs in injured peripheral nerves and found that this senescent SC phenotype have detrimental effects over axonal regeneration in aged or chronically denervated animals, opening novel possibilities to improve functional recovery after peripheral nerves injuries.

Our results demonstrate that senescent Schwann cells arise in peripheral nerves after injury in aging and chronic denervation, and probably correspond to previous observations of SCs exhibiting senescent markers, and populating long acellularized grafts^20, 21^, demonstrating now that these cells appear in uninjured and injured nerves, and are inhibitory for axonal regeneration. The diminished expression of c-Jun on senescent SCs confirm previous works in the context of chronic denervation and aging^19^, but now defining the identity of these low c-Jun expressing Schwann cells. Given that c-Jun is essential for the reprogramming and maintenance of SCs in a repair phenotype^2^, it will be important to define the relationship between c-Jun expression and SC senescence. Previous works have shown that p16 inhibit c-Jun phosphorylation by associating to the N-terminal region of JNK kinases^50^, and that c-Jun suppress p16 expression by binding to his promoter region^36, 37^. As c-Jun expression declines after SCs remains chronically denervated^51^, a transition into a senescent SC phenotype could be associated to upregulation of p16, which in turn could generate a feedforward loop, further preventing c-Jun expression. Indeed, we found that forced overexpression of c-Jun prevents the increase in p16-positive SCs in chronically denervated nerves (Figure 1K, 1L), an intervention that has been shown to rescue the failure in nerve regeneration caused by aging or chronic denervation^19^.

These assumptions are backed by our experiments, as after genetic or pharmacological elimination of senescent cells, the remaining population of non-senescent SC exhibits an increase in c-Jun and a decrease in p16 positivity within the nerve (Figure 4G, 4H and 5F, 5G). This could indicate that the interaction with senescent cells not only negatively effects c-Jun expression in non-senescent SC, but is also a transient effect, that can be reversed or ameliorated by targeted elimination of the population of senescent cells within the nerve.

Taken together, this data points out the plausible dynamics between p16 and c-Jun in sSC, even though as senescence affects a wide range of processes within the cell, further experiments are needed to dissect the specific mechanisms underlying this relationship in SCs.

Our data consistently show the presence of senescent SC in both chronic denervation and aging after nerve transection, although the mechanism associated to the induction of the senescent phenotype is unclear. Injury-induced SC transition into a repair phenotype is associated with re-entry into a replicative state, which is inhibited by regenerating axons invading the distal nerve stump^1^. Nevertheless, in chronically denervated nerves, rSCs continue in a high replicative state, which is one of the triggers of cell senescence^18, 52^. In addition, constant replication is associated to the upregulation of a DNA damage response (DDR), which can also lead to cell senescence^32, 52^. In aged nerves, SC have shown defects in their transcriptional machinery, associated with DNA damage, especially in regeneration-associated genes (RAGs)^53, 54^.

In concordance, senescence markers were not only found in SC but also in SOX10-negative cells, composing up to 60% of the senescent cell population within the nerve. As peripheral nerves are also composed of fibroblasts, perineural and endothelial cells, as well as resident macrophages^55, 56^, these cells might also contribute to the senescent cell population found in damaged nerves in both aging and chronic denervation. Indeed, in both conditions it has been shown that the activity of infiltrating macrophages is characterized by an attenuated phagocytosis and the secretion of proinflammatory factors within the nerve^14, 53, 57^. Importantly, it has been reported that some of these factors, such as iNOS, TNFα, IL-1β, and IL-6, are expressed in both SCs and macrophages^58, 59^. Specifically, high levels of NO have been shown to induce cellular senescence^60^. This raises the question of whether macrophages contribute to SC senescence or become senescent by the SASP released by sSC. Although our experiments *in vitro* demonstrate that siSC-derived SASP directly inhibit axonal growth, secreted molecules from senescent cells other than SCs within the nerve, can also contribute to inhibition of axonal regeneration, or to the induction of cell senescence by a bystander effect, as it has been reported in other damage conditions^61^. Further analysis on the diversity of senescent cells in chronically denervated and aged nerves after injury will be important to fully understand the contribution of different cell populations.

The current hypothesis in the field is that low c-Jun expressing SCs lose their pro-regenerative capacity after chronic denervation and aging^19^. However, our data suggest a more complex scenario, in which sSC acquire a phenotype inhibitory for axonal growth. Although elimination of sSC is sufficient to rescue regenerative capacities in aged and chronically denervated nerves, the appearance of c-Jun expressing Schwann cells after senolysis suggests the possibility that the elimination of the inhibitory sSC component plus the activation of a rSC phenotype is associated with the regenerative response.

Associated with the inhibitory effect of sSC on axonal growth, our *in vitro* data demonstrate an important contribution of secreted factors, which are able to overcome the pro-regenerative effect of secreted factors released by rSCs. Our results demonstrate a high abundance of SASP-related transcripts, proinflammatory cytokines, and proteins in both aged and chronically denervated nerves (Figure 2B and 2C, S2B). These are probably associated with sSC as well as other cells populating the damaged nerve. In addition to SC, is important to consider macrophages, as although they are necessary for regeneration at early stages following nerve injury^62^, their prolonged presence may inhibit regeneration and has been previously associated to tissue dysfunction in aging^16^. Our cytokine array analysis suggests several candidates that might contribute to the inhibitory environment. Interestingly, out of 111 proinflammatory cytokines analyzed, 25 were downregulated in aged or chronically denervated nerves after senolytic treatment, and only 3 showed clear downregulation in both conditions, IL-6, CCL19, and CXCL1. IL-6 is secreted by both SC and macrophages upon injury, and has been associated with several age-related diseases such as multiple sclerosis, Alzheimer disease, diabetes and systemic lupus^63^. Interestingly, macrophages suppress SC proliferation and maturation through IL-6 secretion^64^. Particularly, CXCL1, also secreted by both SC and macrophages, has been associated with physical inhibition of axonal outgrowth in DRG neurons^45^. CXCL1 interferes with the functioning of TRPV1 (Transient Receptor Potential 1) receptors in TRPV1 (+)/IB4 (Isolectin B4) (+) DRG neurons^65^. These effects are important as studies have shown that TRPV1 receptors are physically and functionally present at dynamic neuronal extensions, including growth cones of embryonic and adult DRG neurons^66^. Strikingly, this effect correlates with the growth cone collapse observed in response to conditioned media from siSC (Figure 3F and G). These data demonstrate for the first time that growth cone dynamic is a target of secreted factors from senescent cells, which could have important consequences in other injured tissues that require innervation for proper function.

Taken together our results show that the development of senescence in SCs undermine axonal regeneration, by preventing or counteracting the activation of the reparative program in non-senescent SCs and through direct influence on growth cone dynamics. This effect can be reversed after pharmacological elimination of senescent cells within the nerve, increasing levels of c-Jun and thus the activity of the reparative phenotype of SC. Chronic denervation and aging are the main clinical problems associated to peripheral nerve injuries, even though effective treatment has been elusive due to the incomplete understanding of the underlying causes of this regenerative failure^7^. Our data strongly suggest that SC senescence can be regarded as one of the principal mechanisms associated to inhibition of axonal regeneration. Our approach, using FDA approved drugs currently in clinicals trials for its application as senotherapeutics^67^, effectively broaden the spectrum of its clinical use and effectivity. Finally, this work shows that the pharmacological ablation of the sSC population using a senolytic drug is capable of significantly increasing regeneration performance after nerve injury in aged and chronic denervation conditions, leading us one step closer to improved clinical applications.

## Materials and Methods

### 3.1. Transgenic mice

Mice generated to overexpress c-Jun selectively in Schwann cells as described^68^, conformed to UK Home Office guidelines under the supervision of University College London (UCL) Biological Services under Protocol No. PPL/70/7900. p16-3MR transgenic mice (a C57BL/6J strain) were generated as previously described^69^ (BUCK Institute for Research on Aging, Novato CA), and were bred in the animal facility of the Universidad Mayor.

### 3.2. Tibial nerve transection and reconnection

C57b mice from different ages (adult: 2-4 months-old and aged: 20-22 months-old) were anesthetized with isoflurane (Baxter, Illinois, USA) and the right sciatic nerve was exposed between the hipbone and the sciatic notch. Afterwards, the sciatic branches were isolated, and the tibial nerve was transected and the distal tibial nerve sutured using an 11-0 suture to the nearest muscle to prevent reconnexion. For reconnection surgery, the tibial nerve was detached from the muscle and reconnected with an 10-0 suture to the freshly transected proximal common peroneal nerve (Figure S2A). In all conditions, the nerve was dissected or reconnected 7, 12 or 42-days post injury (dpi), reconnected nerves were dissected 7 days post reconnection, contralateral uninjured nerves used as control (Figure S2A-D). All animal procedures were carried out in accordance to Universidad Mayor Animal Care Committee Guidelines.

### 3.3. Nerve processing and axon regeneration analysis

For β-gal and immunofluorescence (IF) analysis, nerves were fixed by immersion in 4% in PFA. Cellular senescence of fixed nerves was measured by SA-β-gal activity ^26^ and documented under stereomicroscope photography. Later, nerves were embedded in OCT (Sakura Finetek), cryostat sections were cut longitudinally at 20 or 10um thickness and mounted on Superfrost Plus slides (Thermo Fisher Scientific) for IF on confocal microscopy (Leica TCS SP8) against the markers of senescence: histone y-H2AX, H3K9me3, HMGB1, lamin-b1, p16 and P19arf in parallel to SOX-10 SCs marker. Expression analysis was performed using a custom TaqMan® Array FAST plate (Applied Biosystems), specifically against senescence (p16, p21, p51, LaminB1, Trp53bp1) and SCs (c-Jun, Krox-20, mbp) transcripts (see Table 5). RNA extraction and processing were performed according to manufacturer instructions, qPCR was run on a StepOnePlus™ Real-time PCR system.

For axonal regeneration, c-Jun (regeneration) or Stathmin (axon) antibodies were used in parallel to SOX-10 SCs marker (see Table 4 for details on clones and dilutions). Axonal regeneration was quantified from longitudinal sections of reconnected nerves. Fixed immunolabelled 20 μl cryostat sections of reconnected nerves were photographed using confocal microscopy (Leica SP8). Z-stacks images were merge using ImageJ software Collection Stitching tool and analyzed using Oxford’s Imaris software. The reconnection site was recognized and defined in Z-stack images as the region of attachment of the two nerves of different sizes also marked by the 10-0 suture used to reconnect both branches. Starting from the reconnection site, number of regenerating axons and the total nerve width were measured every 200 μm and the combined average ratio between width and axon was graphed using GraphPad Prism Software.

### 3.4. Assessment of sensory function

Pinprick assay was performed as previously described (Ma et al., 2011). An Austerlitz insect pin (size 000) (FST) was gently applied to the plantar surface of the paw without moving the paw or penetrating the skin. The most lateral part of the plantar surface of the hind paw (sensory field of the sciatic nerve) was divided into 5 areas (see Figure S2E). The pinprick was applied (twice) from the most lateral toe (area E) to the heel (area A). A response was considered positive when the animal briskly removed its paw, and the mouse was graded 1 for this area, and then tested for the next one. If none of the applications elicited a positive response, the overall grade was 0. In that case, the saphenous territory of the same paw was tested as a positive control, which in all cases elicited a positive response.

Thermal allodynia was assessed using the Hargreaves apparatus (Ugo Basile, Cat.37370, IT). All measurements were conducted by a blinded observer. Mice were habituated in the Hargreaves apparatus in individual polyvinyl boxes (10×10×14 cm) placed on glass. A radiant light heat source (45 IR) was focused on the plantar skin of the hind paw, and the latency to a withdrawal response was recorded. The mean time to withdrawal was determined from the average of three tests, separated by at least 2 min. A cut-off time of 20 s was used to avoid tissue damage.

### 3.5. Senolysis assays

For senolysis assays, the sciatic nerves of adult (2-4 month-old) and aged (20-22 month-old) C57b mice was sectioned as detailed above. For chronic denervation assays, 28 days after injury, animals were submitted to gavage with a dose of ABT263 (navitoclax) diluted on DMSO and corn oil in a 1:10 proportion respectively (final ABT263 concentration of 50 mg/kg) or vehicle for 5 consecutive days. After treatment, the mice were left for 7 days to allow senolysis to develop. In acute denervation assays, animals were submitted to gavage with a dose of ABT-263 (final ABT263 concentration of 50 mg/kg) or vehicle for 5 consecutive days starting immediately after injury. After treatment, the mice were left for 7 days to allow senolysis to develop. After both treatments, reconnection surgery was performed in the treatment and vehicle groups as described previously. Finally, 7 days after reconnection, animals were sacrificed, and nerves were harvested for immunofluorescence under confocal microscopy as mentioned above.

### 3.6. Schwann cell cultures

SCs were obtained from newborn P2-P3 SD rat sciatic nerves as previously described^70^, expanded over laminin coating and maintained on DMEM 10% FBS supplemented with 2 μM forskolin and 20 μg/ml bovine pituitary extract (BPE). The medium was replaced every 3 days until the cells reached 90% confluence. For senescence induction, cells were seeded at 50,000 cell/cm2 in T175 flasks for exosomes isolation or in 24 well plate for senescence induction and IF.

### 3.7. Senescence induction of Schwann cell cultures

Senescence induction was carried out by adding 80 nM of doxorubicin to SC cultures for 24 hrs. Afterwards, the growth medium was replaced with doxorubicin-free medium and maintained for 9 days in order to achieve senescence induced Schwann cells (siSCs). Cells were fixed on 4% PFA and senescence determination was made by measuring SA-β-gal activity (KAA02 Merck) and IF against the markers of senescence: p16, p21, histone y-H2AX, HMGB1, lamin-b1 and P19arf in parallel to S100 SCs marker (Table 4).

### 3.8. Rat embryonic dorsal root ganglia (DRG) culture

DRG explants were prepared as previously described^71^. Briefly, DRG were dissected from spinal cord of E16.5 rat embryos. DRG were used as explants or reaggregates, explants were seeded in 96-well plates coated with poly-L-lysine/collagen type I and neurobasal medium supplemented with of 1X B27 (50X Gibco), 20 μM L-glutamine, 1X antibiotic-antimycotic solution (100X Gibco), 3.75 μM aphidicolin, 1.25 μM 5-fluoro-2-deoxyuridine and 50 ng/ml of nerve growth factor (NGF-2.5s). For reaggregates, the ganglia were incubated in 0.5% trypsin for 5 min at 37°C, then the tissue was disaggregated and suspended on neurobasal medium supplemented as mentioned above. Later, the disaggregated cells were seeded on 96 wells plates filled with 50ul of 1x agarose solution and agitated on orbital shaker for 15 mins at 37°C then incubated over night at 37°C on C02 chamber. Next day, reaggregated DRGs were carefully transferred using a p200 micro pipette and seeded in 96-well plates coated with poly-L-lysine/collagen type I. After 3 days in culture, DRG explants were documented under transmitted light microscopy (Leica DMi8) and then fixed in 4% PFA, images were quantified for neurite growth area using the ImageJ software, immunofluorescence against medium neurofilament (NF-M) antibody was performed for representative images.

### 3.9. DRG co-culture

For direct interaction DRG reaggregates were co-cultured on 24 wells plates previously seeded with SC or siSCs, after 3 days the co-culture was fixed with 4%PFA for immunofluorescence. Immunofluorescence of co-cultures was performed against NF-M and S100 antibodies for quantification of neurite outgrowth area using the ImageJ software.

### 3.10. Secretome extraction

Conditioned medium (CM) was collected from SCs or siSCs incubated with exosome-depleted FBS/phenol red free SC media, exosomes were depleted as described in ^72^. The media in every condition was extracted and concentrated using 3Kda filter units (Millipore) for 1hr at 5000g. Total protein was quantified by BCA protein assay (Pierce Cat N°23225). For conditioned media assays, 150 µg of concentrated conditioned media protein was added to 250 µl of culture media.

### 3.11. Mass spectrometric protein analysis of Schwann SASP

The conditioned media with 2% FBS of Schwann cells (repair and Doxo-treated cells, n = 3 each) were collected as previously described^40^. Briefly, salt and other media components were removed using 3 kDa cutoff columns (Amicon Centrifugal Filters). Highly abundant proteins, including albumin and IgG, were removed using spin columns (High Select™ Depletion Spin Column). Depleted conditioned media were lysed using lysis buffer (5% SDS and 50 mM TEAB), spun through the micro S-Trap columns (Protifi), and trypsin digested at a 1:25 ratio, 37°C overnight. The peptides were desalted and re-suspended in aqueous 0.2% formic acid for mass spectrometry-based comparative quantification analysis. Mass Spectrometric Data Independent Acquisition (DIA)^73, 74^ was performed on a Dionex UltiMate 3000 system coupled to an Orbitrap Eclipse Tribrid mass spectrometer (Thermo Fisher Scientific, San Jose, CA); (detailed parameters provided as supplementary information). DIA data was processed in Spectronaut v15 (version 15.1.210713.50606; Biognosys) using directDIA. Identification was performed using a 1% precursor and protein q-value. Quantification was based on the MS2 area, local normalization was applied, and iRT profiling was selected. Differential protein expression analysis was performed using a paired t-test, and q-values were corrected for multiple testing, specifically applying group-wise testing corrections using the Storey method^75^.

### 3.12. Transcriptomic analysis of denervated nerves

Two month old adult Sprague-Dawley rats underwent sciatic nerve transection at the upper thigh as described previously^76^. The proximal sciatic nerve was sutured to nearby muscles to prevent reinnervation of the distal denervated sciatic nerve. At different time points from 1 day to 180 days, the distal denervated and contralateral control uninjured sciatic nerves were harvested and frozen on dry ice for microarray analysis as described previously^77^. Total RNA was extracted, RNA quantity was assessed with Nanodrop Spectrophotometer (Nanodrop Technologies) and quality with the Agilent Bioanalyzer (Agilent Technologies). As per the manufacturer’s protocol, 200 ng of total RNA were amplified, biotinylated and hybridized to Illumina Rat Expression Beadchips, querying the expression of 22,000 RefSeq transcripts. Four replicates were run per sample category. Raw data were analyzed by using Bioconductor packages and contrast analysis of differential expression was performed using the LIMMA package^78^. After QC and linear model fitting, a Bayesian estimate of differential expression was calculated and the false discovery rate was set at 5% all as per standard protocols^79^.

Differentially expressed genes were contrasted against SASP-related genes database, using the SASP query tool (www.saspatlas.com).

### 3.13. Cytokine array

Aged or chronically denervated nerves treated with ABT263 or vehicle were extracted as described above. An inflammatory cytokines array (RD.ARY028, R&D Systems) was used according to manufacturer’s instructions to analyze actual concentration of SASP inflammatory proteins found in our analysis.

## 4. Data availability

Raw data for all the images used in this work are available in the EBI BioStudies database (https://www.ebi.ac.uk/biostudies/) with the accession number TMP_1679064586359

## Acknowledgments and funding

This work was supported by grants from the Geroscience Center for Brain Health and Metabolism, FONDAP-15150012, Fondo Nacional de Desarrollo Científico y Tecnológico (FONDECYT) N° 1150766, Michael J Fox Foundation for Parkinson’s Research 17303 (F.A.C.). Dr. Miriam and Sheldon G. Adelson Medical Research Foundation (A.H. and D.H.G.). National Institutes of Health under award numbers U01 AG060906 (B.S.), P01 AG017242 and R01 AG051729 (J.C.). We acknowledge the support of instrumentation for the Orbitrap Eclipse Tribrid from the NCRR shared instrumentation grant 1S10 OD028654 (B.S.). S.K.P. is supported by a Glenn Fellowship in Aging Research from the Glenn Foundation.

