## Supplementary figures and images for "Senescent Schwann cells induced by aging and chronic denervation impair axonal regeneration after peripheral nerve injury"

### Figure S1

Supplementary Figure 1

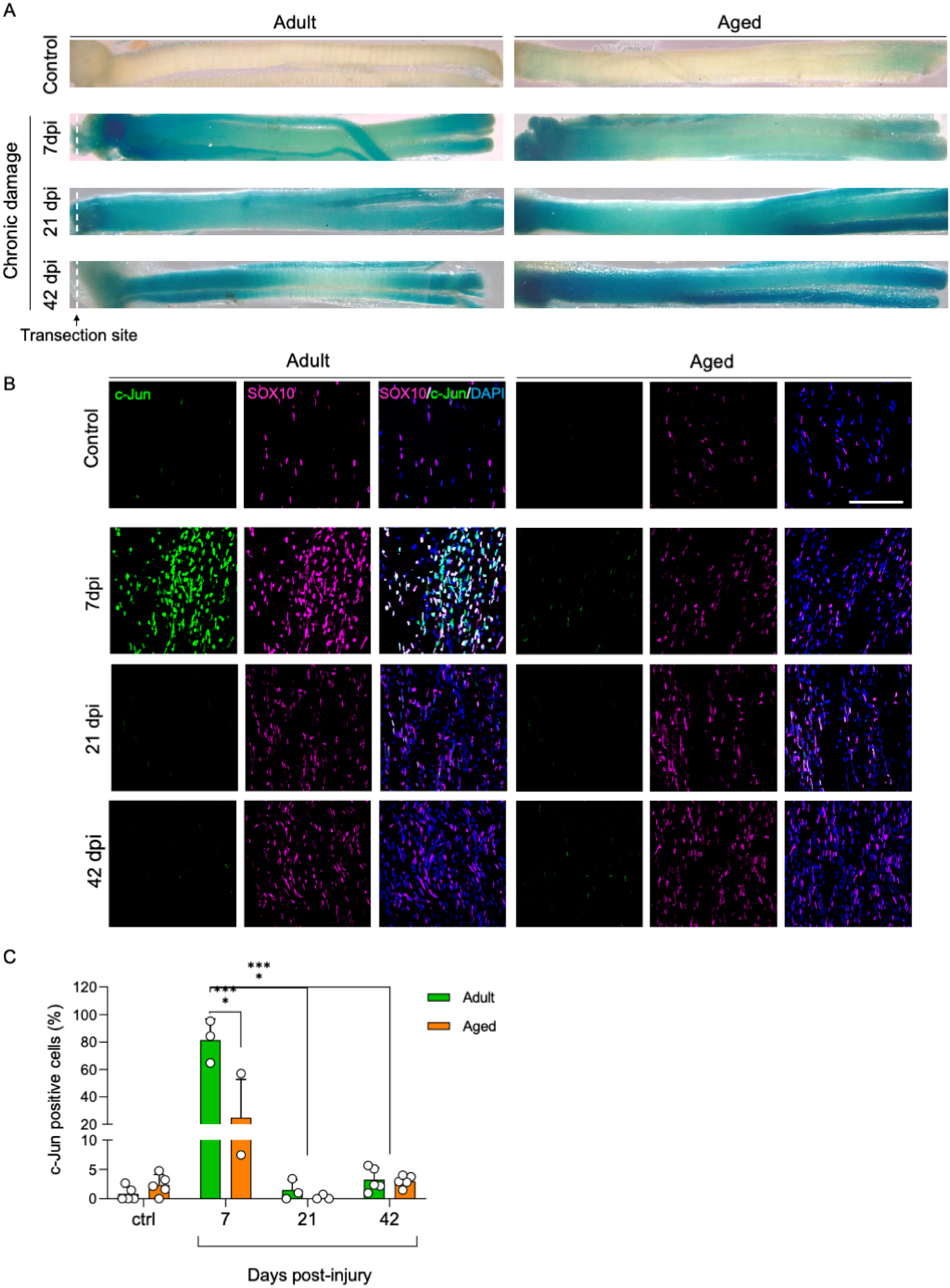

### Figure S2

Supplementary Figure 2

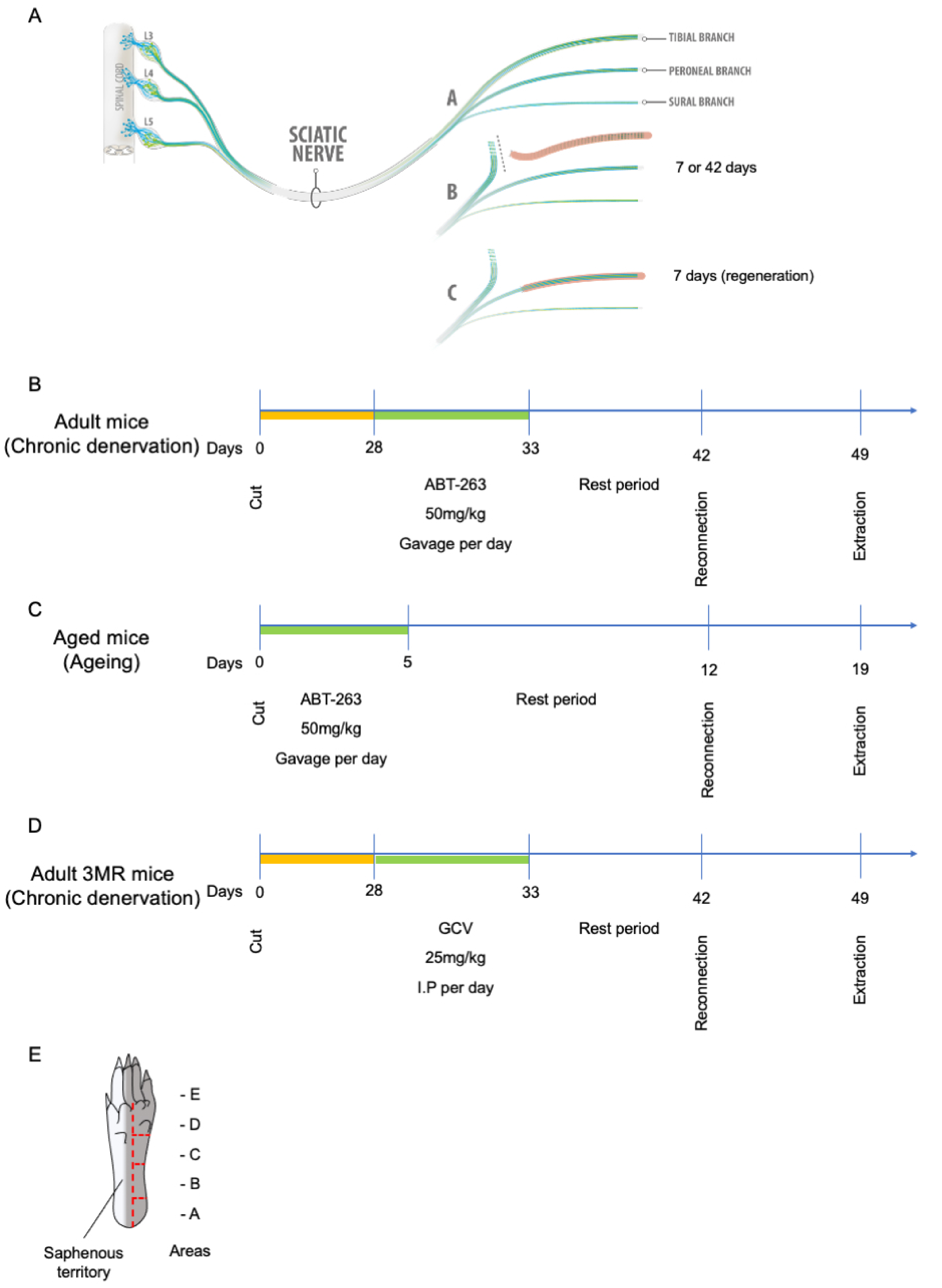

### Figure S4

Supplementary Figure 4

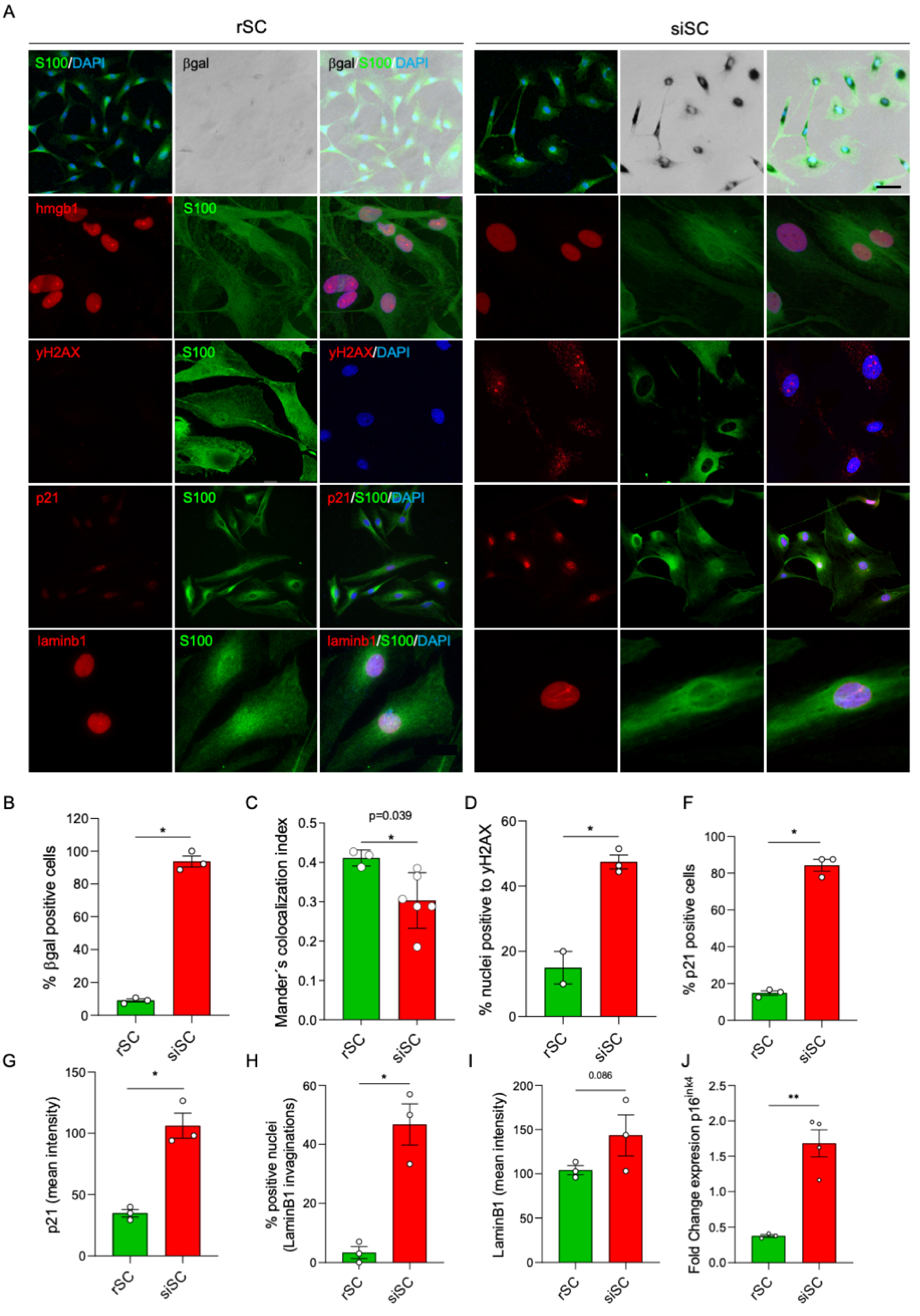

### Figure S5

Supplementary Figure 5

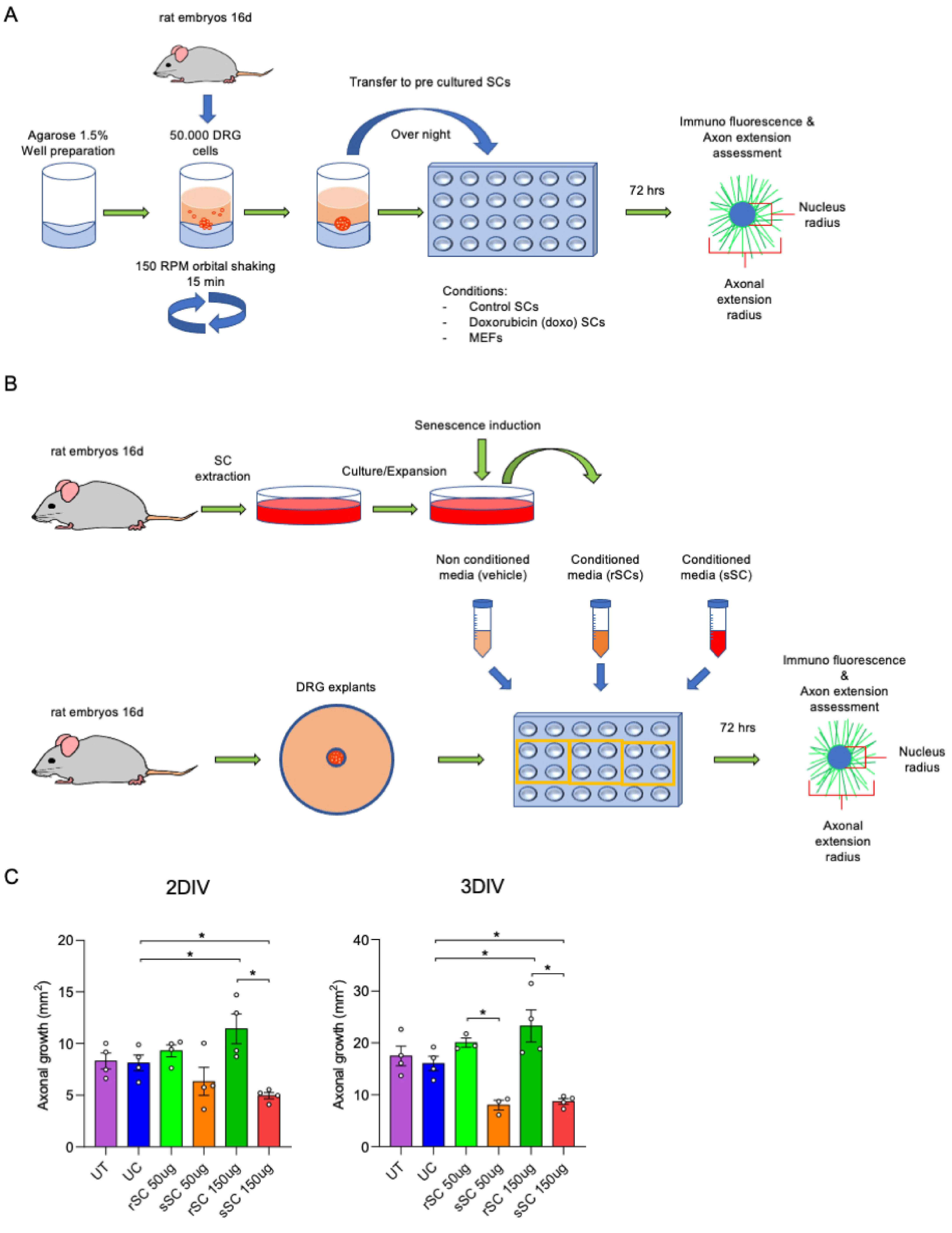
