## Supplementary material for "Senescent Schwann cells induced by aging and chronic denervation impair axonal regeneration after peripheral nerve injury": Figure S3

Supplementary Figure 3

A

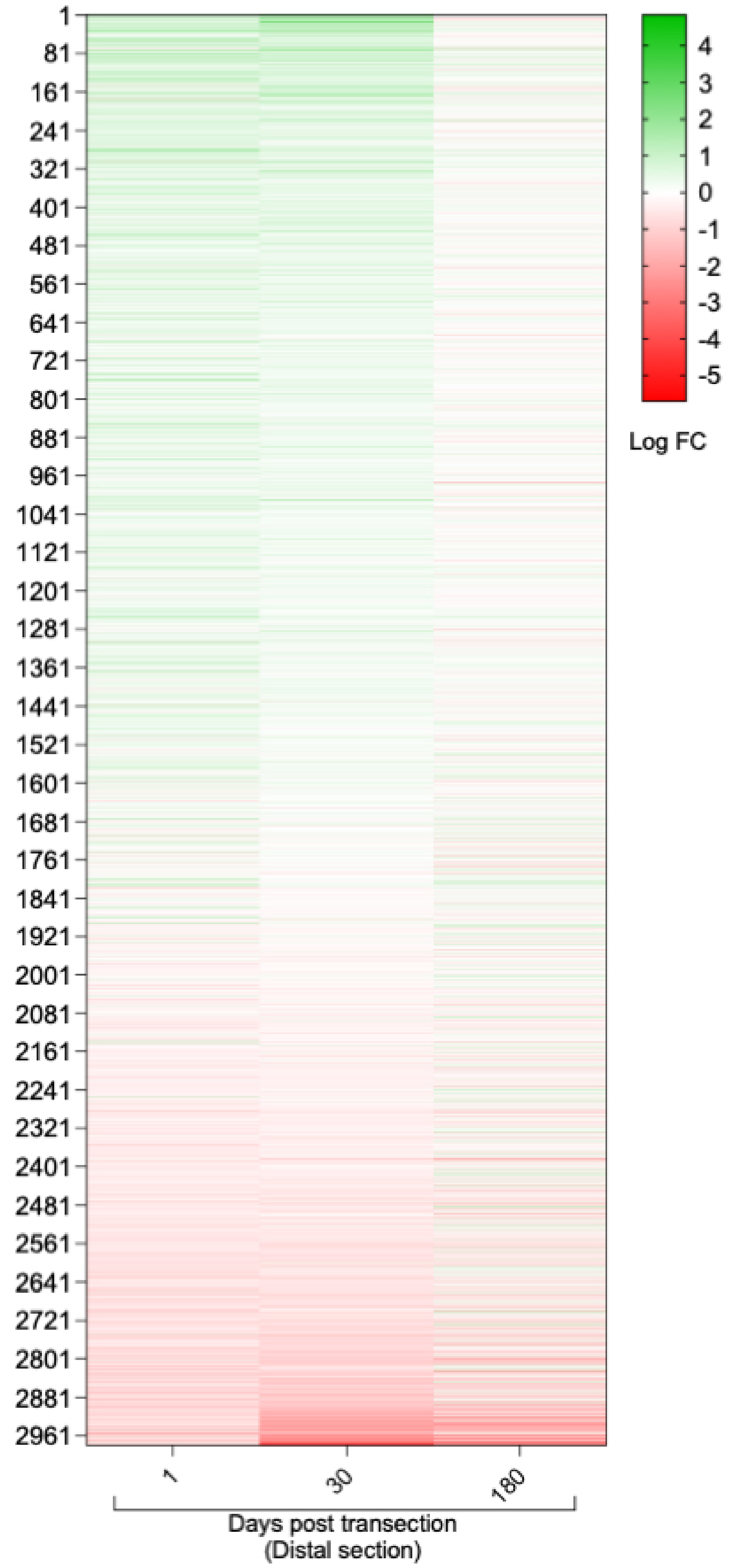


B

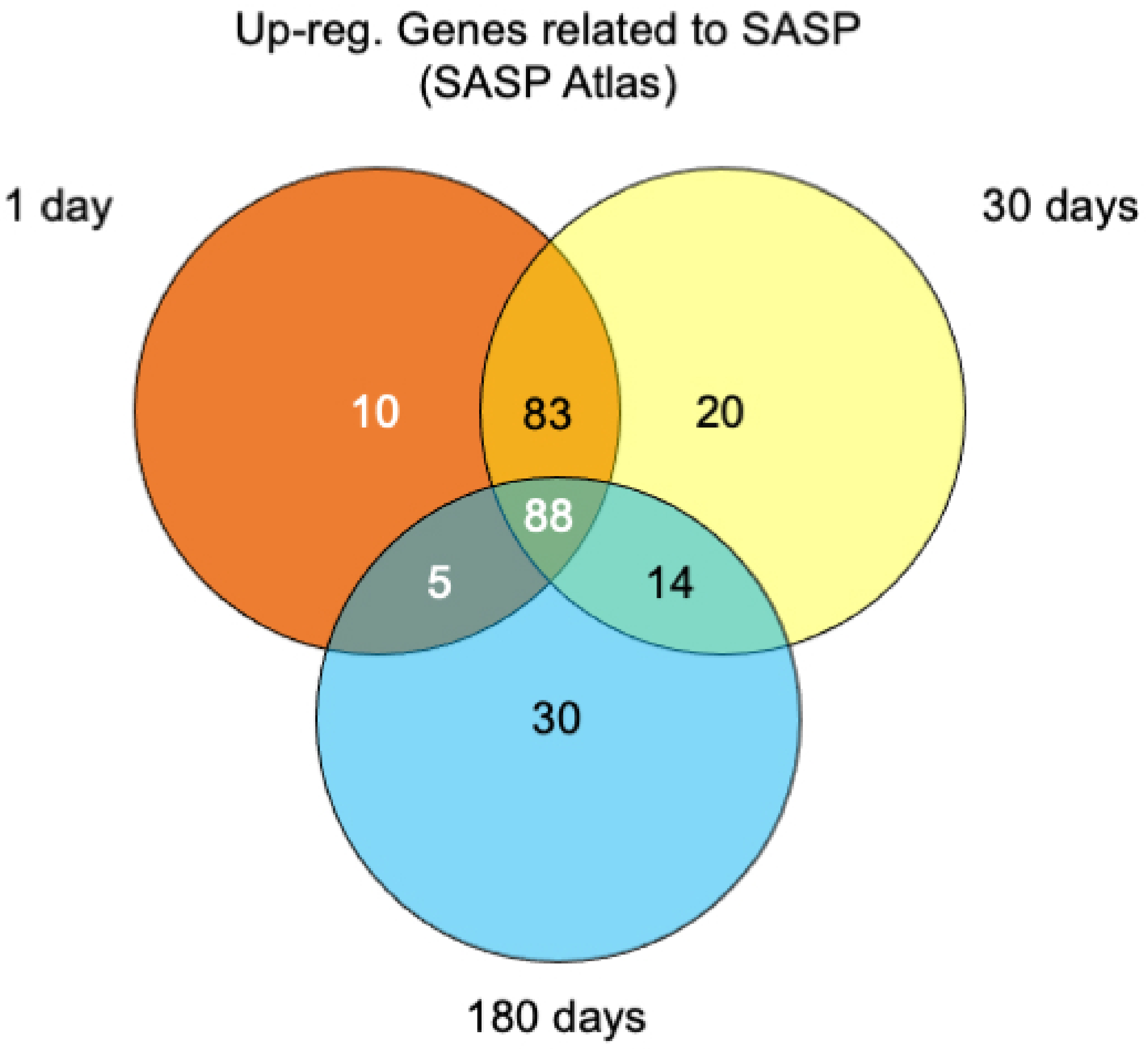


C

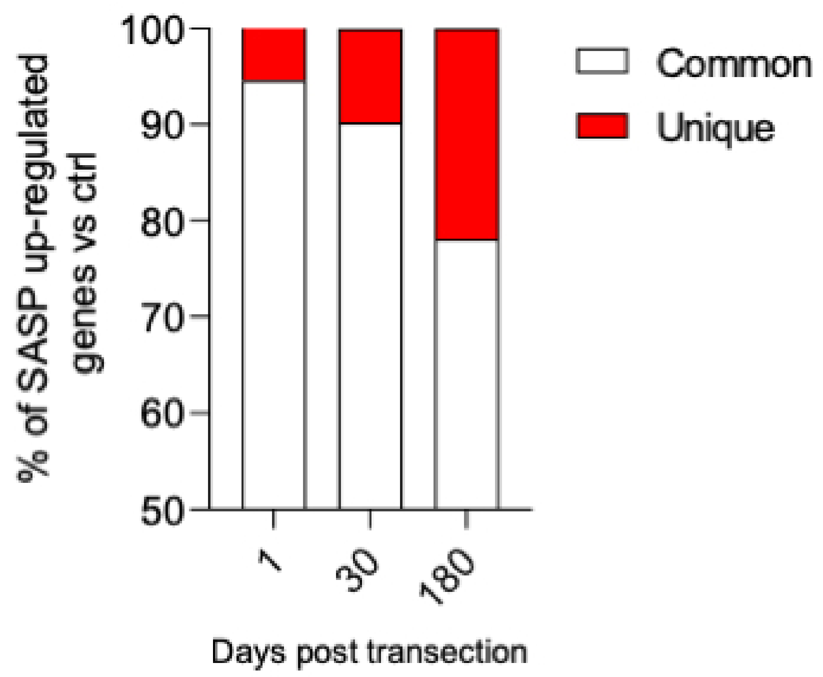


D

| Unique SASP components | | |
|---|---|---|
| 1 day | 30 days | 180 days |
| ARCN1 | PSMA2 | SOD3 |
| CCT8 | CBX5 | ITIH3 |
| ATIC | TXNL1 | MRPS5 |
| PRDX1 | SERPING1 | HSPB1 |
| CSTB | SNX6 | AOC3 |
| TALDO1 | GAS6 | ADD3 |
| MIF | PIGK | EPS8L2 |
| RAB5C | FAP | FBLN1 |
| MVD | PCOLCE | ENPP2 |
| PSMB3 | IMPA1 | MAMDC2 |
| | HEXA | SF3B1 |
| | CASK | ACADM |
| | MYH10 | CD59 |
| | HEBP2 | PGRMC1 |
| | UCHL1 | ECH1 |
| | SDF4 | ITM2B |
| | NPC2 | EPRS |
| | HEXB | MFGE8 |
| | FBLN5 | CST3 |
| | LGALS1 | PC |
| | | EIF4A2 |
| | | KPNA1 |
| | | TRIP10 |
| | | GSTT1 |
| | | RNASE4 |
| | | SDHA |
| | | APP |
| | | MDH1 |
| | | GRHPR |
| | | MPST |
