## Supplementary Figure legends for "Senescent Schwann cells induced by aging and chronic denervation impair axonal regeneration after peripheral nerve injury"

**Figure S1. Sciatic nerves from chronically denervated and aged animals have diminished c-jun expression and axonal regeneration in SCs after injury compared to adult animals.** (A) Representative brightfield images of β-galactosidase activity on non-injured and chronically denervated (42dpi) whole sciatic nerves of adult and aged animals after b-galactosidase assay. Scale bar, 1000 μM. (B, C) Representative IF confocal images (SOX10, magenta; c-Jun, red; DAPI, blue) and graph comparison of c-Jun activation in the nucleus of SCs on longitudinal cryostat sections of adult and aged mice sciatic nerves 7, 21 and 42 dpi. Scale bar, 100 μM.

**Figure S2. Schematic representation of *in vitro* experiments.** (A) Schematics of the co-culture protocol. (B) Schematics of the conditioned media treatment protocol. (C) Comparison between DRG re-aggregates after 2 or 3 days in vitro (DIV) of exposure to unconditioned media (UM) and conditions media from SC or sSCs in concentrations of 50 or 150ug of concentrated protein from collected media (UT, untreated DRGs). (n=3-4 re-aggregates per group; * p<0.05 by Student’s t-test compared between conditions; error bars indicate S.D.).

**Figure S3. Differentially expressed genes after chronic denervation involved in the senescence-associated secretory phenotype (SASP).** (A) Differentially expressed genes in adult distal nerves after 1, 30 and 180 days after sciatic nerve transection. (B) Venn diagram of up-regulated genes after 1, 30 and 180 days after transection contrasted against the SASP-ATLAS database. (C-D) Graph comparison of percentage of unique SASP genes against common SASP genes after 1-, 30- or 180-days post transection and table of unique SASP genes in every condition.

**Figure S4. Markers of senescence in primary Schwann cell culture after doxorubicin treatment**. (A) Representative IF images in rSC and siSC, S100, green, B-gal, black; Hmgb-1/yH2AX, red; DAPI, blue. Scale bars, βgal, 100 μM; hmbgb-1, 25 μM; yH2AX, 50 μM; p21, 10 μM; laminb1, 25 μM. (B-J) Graph comparison of β-gal^+^ cells, mander´s co-localization index of HMGB1, yH2AX foci/nucleus (D), p21 positive cells (F) and expression levels (G), nuclei positive for LaminB1 marked invaginations (H) and expression (mean intensity) (I), p16INK4a fold change (qRT-PCR) (J) between non-senescent and siSCs and rSCs (*p<0.05 by Student’s t-test compared between conditions; error bars indicate S.D.).

**Figure S5. Schematic representation of *in vivo* experiments.** (A-D) Tibial nerve was transected and sutured to the muscle. After 28 days for adults and 5 days in aged animals, ABT, GCV or vehicle treatment was performed for 5 consecutive days. After 40 dpi in adult or 12 dpi in aged, nerves were sutured to freshly transected peroneal nerve. Then, 7days after reconnection, nerves were harvested for senescent and regeneration markers analysis (b-gal, IF or qPCR). (E) Schematic representation of pinprick touch points.
