## Supplementary methods for "Senescent Schwann cells induced by aging and chronic denervation impair axonal regeneration after peripheral nerve injury"

**Mass spectrometric protein analysis of Schwann cells senescence-associated secretory phenotype (SASP) and statistical data processing**

The conditioned media with 2% FBS from Schwann cells (repair and Doxo-treated cells [n = 3 each]) were collected as previously described^1^. Salt and other media components were removed using 3 kDa cutoff columns (Amicon Centrifugal Filters). Highly abundant proteins, including albumin and IgG, were removed using spin columns (High Select™ Depletion Spin Column). Depleted conditioned media were lysed using lysis buffer (5% SDS and 50 mM TEAB). Each extract was reduced by incubation with 20 mM dithiothreitol in 50 mM TEAB for 10 min at 50ºC and subsequently alkylated with 40 mM iodoacetamide in 50 mM TEAB for 30 min at RT in the dark. Extracts were acidified to pH < 1 using phosphoric acid (v/v), and 100 mM TEAB in 90% methanol was added. The protein extract was spun through the micro S-Trap columns (Protifi), washed with 90% methanol in 100 mM TEAB, and then placed in clean elution tubes for trypsin digestion at a 1:25 ratio in trypsin digestion buffer (50 mM TEAB, pH ~8) (protease: protein, wt:wt) overnight. Peptides were then sequentially eluted with 50 mM TEAB and 0.5% formic acid and 50% acetonitrile in 0.5% formic acid. Both fractions were pooled together, vacuum-dried, and re-suspended in 0.2% formic acid for desalting. The desalted peptides were concentrated and re-suspended in aqueous 0.2% formic acid for mass spectrometry-based quantitative analysis.

**Mass Spectrometric Data Independent Acquisition (DIA)**

LC-MS/MS analyses were performed on a Dionex UltiMate 3000 system coupled to an Orbitrap Eclipse Tribrid mass spectrometer (Thermo Fisher Scientific, San Jose, CA). The solvent system consisted of 2% ACN, 0.1% FA in H2O (solvent A), and 98% ACN, 0.1% FA in H2O (solvent B). Proteolytic peptides (50 ng) were loaded onto an Acclaim PepMap 100 C18 trap column (0.1 x 20 mm, 5 µm particle size; Thermo Fisher Scientific) for 5 min at 5 µL/min with 100% solvent A. Peptides were eluted on an Acclaim PepMap 100 C18 analytical column (75 µm x 50 cm, 3 µm particle size; Thermo Fisher Scientific) at 300 nL/min using the following gradient of solvent B: 2% for 5 min, linear from 2% to 20% in 125 min, linear from 20% to 32% in 40 min, up to 80% in 1 min, 80% for 9 min, down to 2% in 1 min, and 2% for 29 min, for a total gradient length of 210 min.

Schwann cells SASP peptides described above were acquired in DIA mode. Full MS spectra were collected at 120,000 resolution (AGC target: 3e6 ions, maximum injection time: 60 ms, 350-1,650 m/z), and MS2 spectra at 30,000 resolution (AGC target: 3e6 ions, maximum injection time: Auto, NCE: 27, fixed first mass 200 m/z). The isolation scheme consisted of 26 variable windows covering the 350-1,650 m/z range with an overlap of 1 m/z (Table S2)^2^.

**DIA data processing and statistical analysis**

DIA data was processed in Spectronaut v15 (version 15.1.210713.50606; Biognosys) using directDIA. Acquired data were searched against the *homo sapiens* proteome with 42,789 protein entries (UniProtKB-TrEMBL), accessed on 12/07/2021. Trypsin/P was set as a digestion enzyme, and two missed cleavages were allowed. Cysteine carbamidomethylation was selected as a fixed modification, while methionine oxidation and protein N-terminus acetylation as variable modifications. The data extraction parameter was set as dynamic. Identification was performed using a 1% precursor and protein q-value (experiment). Quantification was based on the MS2 area, local normalization was applied, and iRT profiling was selected. Differential protein expression analysis was performed using a paired t-test, and q-values were corrected for multiple testing, specifically applying group-wise testing corrections using the Storey method^3^. Table S1 enlists protein groups with at least one unique peptide, q-value < 0.05, and absolute Log2 (fold-change) > 0.58 that are considered significantly altered.

1.	Basisty, N. *et al.* A proteomic atlas of senescence-associated secretomes for aging biomarker development. *PLoS Biol.* **18**, e3000599 (2020).

2.	Bruderer, R. *et al.* Optimization of experimental parameters in data-independent mass spectrometry significantly increases depth and reproducibility of results. *Mol. Cell. Proteomics* **16**, 2296–2309 (2017).

3.	Burger, T. Gentle Introduction to the Statistical Foundations of False Discovery Rate in Quantitative Proteomics. *J. Proteome Res.* **17**, 12–22 (2018).
